# Comparative molecular and immunoregulatory analysis of extracellular vesicles from *Candida albicans* and *Candida auris*

**DOI:** 10.1101/2020.11.04.368472

**Authors:** Daniel Zamith-Miranda, Heino M. Heyman, Sneha P. Couvillion, Radames J. B. Cordero, Macio L. Rodrigues, Leonardo Nimrichter, Arturo Casadevall, Rafaela F. Amatuzzi, Lysangela R. Alves, Ernesto S. Nakayasu, Joshua D. Nosanchuk

## Abstract

*Candida auris* is a recently described multidrug-resistant pathogenic fungus that is increasingly responsible for healthcare associated outbreaks across the world. Bloodstream infections of this fungus cause death in up to 70% of the cases. Aggravating this scenario, *C. auris’* disease-promoting mechanisms are poorly understood. Fungi release extracellular vesicles (EVs) carrying a broad range of molecules including proteins, lipids, carbohydrates, pigments, and RNA, many of which are virulence factors. Here, we carried out a comparative molecular characterization of *C. auris* and *C. albicans* EVs and evaluated their capacity to modulate effector mechanisms of host immune defense. Using proteomics, lipidomics, and transcriptomics, we found that *C. auris* released EVs with payloads that were strikingly different from EVs released by *C. albicans*. EVs released by *C. auris* potentiated the adhesion of this yeast to an epithelial cell monolayer. *C. auris* EVs also induced the expression of surface activation markers and cytokines by bone marrow-derived dendritic cells. Altogether, our findings show distinct profiles and properties of EVs released by *C. auris* and by *C. albicans,* and highlight the potential contribution of *C. auris* EVs to the pathogenesis of this emerging pathogen.

## Introduction

*Candida auris* is a recently described pathogenic fungus that has emerged as a serious cause of healthcare associated infections across the world ^1^. Therefore, it is considered a global threat by the US Center for Disease Control and Prevention ^2^. The biological challenges for combatting *C. auris* include the fungus’ capacity to form resilient biofilms and to resist multiple antifungal drugs ^3^. *C. auris* kills 30-70% of the infected individuals ^4^. Although we have deep knowledge regarding the disease-promoting mechanisms deployed by other *Candida* species, relatively little is known about *C. auris*. We have recently compared the molecular profiles of two *C. auris* isolates vs. *Candida albicans* by integrating proteins, lipids, and metabolites of these yeast cells, and demonstrated that *C. auris* has an elevated expression of pathways related to drug resistance and virulence, such as sterol metabolism and drug resistance-related transporters ^5^.

Disease development is a combination of fungal virulence factors and the affected host’s ability to efficiently control the fungal growth, and extracellular vesicles (EVs) play a role in both of these factors. EVs are lipid bilayered structures released by a broad variety of uni- or multicellular organisms ^6^. Fungal EVs from *Cryptococcus neoformans* were first described in 2007 ^7^ and they have since been shown as an important mechanism for molecular export in a variety of fungal species. EVs produced by fungi carry many biologically active molecules, including virulence factors and regulators, indicating that they could activate the innate immune system and influence disease development ^8–16^. *In vitro,* fungal EVs impact phagocyte activity, promoting an increase in cytokine levels, modulating phagocytosis and regulating macrophage polarization ^8–10,14,16,17^. Together, these data strongly suggest that fungal EVs activate the immune response. Indeed, *Galleria mellonella* larvae are protected by pre-treatment with EVs from *C. albicans, C. neoformans* and *Aspergillus flavus*^9,18,19^. Recently, we demonstrated that immunization of mice with EVs from *C. albicans* confers full protection against systemic candidiasis ^11^.

However, the outcome of fungal EV and host response depends on the model investigated. For instance, yeast EVs released from co-cultures of dendritic cells (DCs) and *Malassezia sympodialis* induce the production of TNF-α and higher levels of IL-4 by PBMC from patients with atopic eczema, when compared to control PBMC, displaying an allergic reaction ^20,21^. *C. neoformans* and *Sporothrix brasiliensis* EVs are associated with virulence and disease progress in murine models, respectively ^22,23^. We hypothesize that the multiple activities attributed to fungal EVs could be dependent on their composition, which at least partially differs according to the species investigated ^9,21,24–29^. Thus, a more complete analysis on EVs composition could open new views for understanding fungal diseases.

Here, we performed a detailed characterization of EVs released by *C. auris* and *C. albicans.* Differences in size and sterol/protein ratios were observed. Using integrated multi-omics (proteomics, lipidomics and, transcriptomics) analysis we compared EVs and whole cells of *C. auris* and *C. albicans* and demonstrated significant compositional differences that could impact pathogenesis. Developing functional assays, we demonstrated that *C. auris* EVs influence adhesion to epithelial cells and activation of dendritic cells. Together our results show that *C. auris* produces EVs with a distinct composition in comparison with *C. albicans*, and *C. auris* EVs modulate host cell defense mechanisms.

## Methods

### Cell lines

Two well characterized *C. auris* clinical isolates (MMC1 and MMC2) were acquired from Montefiore Medical Center (NY, USA) ^5^. *C. albicans* strain (ATCC #90028), RAW 264.7 macrophages (ATTC #TIB-71) and HeLa cells (ATTC #CCL-2) were obtained from ATCC. Yeast cells were cultivated in YPD broth and seeded onto Sabouraud agar plates. For each experiment, colonies were inoculated in Sabouraud broth for 24 h at 30 °C before use. J774 and HeLa cell lines were cultivated up to the 10^th^ passage in DMEM supplemented with 10% FBS and 1% non-essential amino acids.

### EVs isolation

One colony of each strain of *C. auris* or *C. albicans* was inoculated in 10 mL of Sabouraud broth for 24 h at 30 °C, and then expanded in 200 mL of fresh medium. After an additional 24 h at 37 °C, the cells were centrifuged. The supernatant was filtered and concentrated 40 fold using an Amicon system with a 100-KDa molecular weight cutoff membrane. The concentrate was then centrifuged twice at 150.000 × *g,* with a PBS washing step between centrifugations. The EVs pellet was suspended in filtered PBS for most of the experiments, and in 50 mM ammonium bicarbonate for proteomic and lipidomic analyses.

### Transmission Electron Microscopy

EVs pellets were fixed in 2.5% glutaraldehyde and 3 mM MgCl2 in 0.1 M sodium cacodylate buffer, pH 7.2 overnight at 4 °C. Samples were then rinsed with buffer and post-fixed in 0.8% potassium ferrocyanide reduced 1% osmium tetroxide in the buffer for 1 h on ice in the dark. After a 0.1 M sodium cacodylate buffer rinse, the samples were incubated at 4 °C overnight in the same buffer. Samples were rinsed with 0.1 M maleate buffer, *en bloc* stained with 2% uranyl acetate (0.22 μm filtered, 1 h, dark) in 0.1 M maleate, dehydrated in a graded series of ethanol and embedded in Eponate 12 (Ted PElla) resin. Samples were polymerized at 37 °C for 2 days and at 60 °C overnight. Thin sections, 60 to 90 nm, were cut with a diamond knife on a Reichert-Jung Ultracut E ultramicrotome and picked up with formvar coated copper slot grids. Grids were stained with 2% uranyl acetate in 50% methanol, followed by lead citrate, and observed with a Phillips CM120 transmission electron microscope at 80 kV. Images were captured with an AMT XR80 high-resolution (16-bit) 8 Mpixel camera.

### Protein and ergosterol quantification

Protein and sterols were quantified using BCA Protein Assay (Thermo) and Amplex Red Cholesterol Assay (Thermo) kits, respectively. Both contents were expressed as a function of the number of yeast cells present in each culture at harvest time.

### Hydrodynamic size distribution of extracellular vesicles by Dynamic light scattering

EVs were suspended in PBS and their hydrodynamic size distributions were measured in a BI-90 Plus Particle Size Analyzer (Brookhaven Instruments) at room temperature as described ^30^. Vesicle preparations were first centrifuged at 13,000 rpm for 5 minutes to remove any larger particles and aggregates. One hundred microliters of sample were loaded into disposable cuvette (Eppendorf 952010077) and analyzed by DLS. The average size distribution was calculated from duplicates of ten individual measurements.

### Isolation and sequencing of extracellular vesicles RNAs

The RNA molecules were isolated with the miRNeasy mini kit (Qiagen) according to the manufacturer’s protocol, to obtain small RNA-enriched fractions. The sRNA profile was assessed in an Agilent 2100 Bioanalyzer (Agilent Technologies). The purified sRNA, from three independent biological replicates, was used for RNA-seq library construction using TruSeq small RNA kit according to the manufacturer’s recommendations. The sequencing was performed with an Illumina HiSeq 2500 platform, TruSeq SBS Kit v3-HS 50 cycles kit (Illumina).

### In silico data analysis

The RNA-seq analysis was performed with CLC Genomics Workbench© software version 12. The reference genomes used for mapping were obtained from the NCBI database (GCA_002775015.1 Cand_auris_B11221_V1). The alignment was performed as follows: additional 100-base upstream and downstream sequences; 10 minimum number of reads; 2 maximum number of mismatches; −2 nonspecific match limit, and minimum fraction length of 0.8 for the RNA mapping. The minimum reads similarity mapped on the reference genome was 80%. Only uniquely mapped reads were considered in the analysis. The libraries were normalized per million and the expression values for the transcripts were registered in TPM (Transcripts per Million). The statistical test applied was the DGE (Differential Gene Expression). For the ncRNA the database used was the ncRNA from the *Candida* genome database: C_auris_B8441_version_sXX-mYY-rZZ_other_features_no_introns.fasta.gz. Gene Ontology analysis was performed using the DAVID annotation tool ^31^.

### Lipidomics and proteomics analyses of extracellular vesicles

Sample processing and analysis were carried out as described^32,33^. Briefly, samples were submitted to simultaneous Metabolite, Protein and Lipid Extraction (MPLEx) ^34^. Extracted lipids were dried in a vacuum centrifuge and dissolved in methanol before analysis by liquid chromatography-tandem mass spectrometry (LC-MS/MS) on a Velos Orbitrap mass spectrometer (Thermo Fisher). Lipid species were identified and manually inspected for validation using LIQUID ^35^. The intensities of each lipid species were extracted using MZmine 2.0 ^36^. Proteins were reduced with dithiothreitol, alkylated with iodoacetamide and digested with trypsin. Resulting peptides were analyzed by LC-MS/MS on a Q-Exactive mass spectrometer (Thermo Fisher). Peptides were identified using MaxQuant (v.1.5.5.1) ^37^ by searching against the *C. albicans* SC5314 and *C. auris* sequences from Uniprot Knowledge Base (downloaded December 6, 2017). Intensity-based absolute quantification (iBAQ) method was used for quantification ^38^. The iBAQ values for individual proteins were normalized against the total sum of all proteins, resulting in the relative protein copy number (percentage from total). *C. auris* and *C. albicans* proteins were considered orthologs with ≥ 40% amino acid sequence similarity ^39^. Heatmap and clustering were performed with MultiExperiment Viewer (MeV) ^40^ or R software and Complex Heatmap package ^41^. For calculating fold changes and plotting the heatmaps, missing values were filled with half of the minimum value of the dataset. Function-enrichment analysis was done with DAVID ^42^, using default parameters.

### Adhesion assay to epithelial monolayers

HeLa cells were seeded on coverslips placed in 24-wells plates and incubated for 24 h at 37 °C. Cell monolayers were preincubated with *C. auris* or *C. albicans* EVs (10 μg/mL of protein) for 1 h and challenged with respective yeasts (pre-stained with NHS-Rhodamine for 30 minutes at 30 °C under shaking) for 1 h in a ratio of 20 yeast cells per HeLa cell. NHS-Rhodamine staining does not change yeast cell growth rates or other cellular characteristics (data not shown). After extensive washing with PBS to remove non-adherent yeast, the cells were fixed with formalin and mounted with mounting media containing DAPI. Images were taken using a fluorescence microscope (Zeiss Imager Z1) and the adhesion was measured by the ratio between NHS-Rhodamine-positive cells divided by DAPI-positive cells for each field, using ImageJ. At least 8 fields containing approximately 400 epithelial cells per field from each slide were counted.

### Analysis of bone marrow-derived dendritic cells (BMDC) activation by extracellular vesicles

BMDC were differentiated as described ^43^. Briefly, bone marrow cells were isolated from male C57BL/6 mice (approved protocol #2014-0501) by flushing both tibias and femurs with RPMI supplemented with 10% of fetal bovine serum (FBS). Bone marrow cells were cultivated for 10 days at 37 °C in the presence of GM-CSF (Peprotech) ^43^. Cultures were fed with media containing GM-CSF at days 3, 6 and 8. BMDC phenotype was evaluated on day 10 by the surface exposure of CD11c and MHCII. At day 10 of differentiation, BMDC were incubated with 1 and 10 μg/mL (protein) of EVs from *C. auris* and *C. albicans* for 24 h at 37 °C and 5% CO2. After this period, cytokines IL-6, IL-10, IL-12p70, TNF-α, and TGF-β were measured in the culture supernatants using ELISA. BMDC were labeled with antibodies (α-CD11c, α-MHCII, α-CD80 and, α-CD86) to evaluate their purity and activation state using flow cytometry.

### Modulation of effector functions of macrophages by extracellular vesicles

**Phagocytosis** – RAW 264.7 macrophages were plated onto 96-well plates and incubated for 24 h at 37 °C. Cells were then incubated with EVs from *C. albicans* or *C. auris* (10 μg/mL of protein) for 1 h until challenge with the respective yeast cell at 1:2 (macrophage : yeast) for 1 h. Plates were washed to remove extracellular yeast cells and then lysed with sterile water for CFU analysis. **Killing**-Bone marrow cells were harvested from C57BL/6 mice as detailed above and incubated with RPMI medium containing 10% of fetal bovine serum and 20% of L929 supernatant at 37 °C. On the fourth day, new medium containing L929 supernatant was added to the culture. On the seventh day of cultures, the cells had matured to differentiated macrophages, confirmed by the expression of F4/80 and absence of LY6C. BMDM were plated in 96-well plates and incubated at 37 °C for 24 h. Cells were incubated with EVs for 4 h at 37 °C until the challenge with yeast cells at 10:1 (macrophage:yeast) for 24 h at 37 °C. Cells were lysed and the suspensions plated onto Sabouraud plates for CFU counting.

### Statistical analyses

All experiments were performed at least 3 independent times, unless stated otherwise. Data sets were analyzed using One-way ANOVA, and Dunnett multi comparison post-test using GraphPad Prism 8. All *p* values lower than 0.05 were considered significant.

## Results

### Morphological characterization of *C. auris* extracellular vesicles

EVs were isolated from the supernatant of *C. albicans* and *C. auris* and then analyzed by transmission electron microscopy (TEM). As reported previously, EVs from *C. albicans* are round and bilayered particles (Figure 1A) ^9^. Similar results were observed for EVs from both *C. auris* isolates (Figure 1B and 1C), consistent with the reported morphology of other fungal EVs ^7,9,12,16,20,23,25,44^. EVs were also analyzed by dynamic light scattering (DLS) to evaluate their global size. The size of EVs isolated from *C. albicans* and *C. auris* MMC2 were very similar, ranging from 50 and 70 nm and the second population between 170-250 nm (Figure 1A and 1C). *C. auris* MMC1 produced EVs of larger hydrodynamic size, ranging from 100 to 150 and the second population between 280 and 370 nm (Figure 1B).

**Figure 1.**
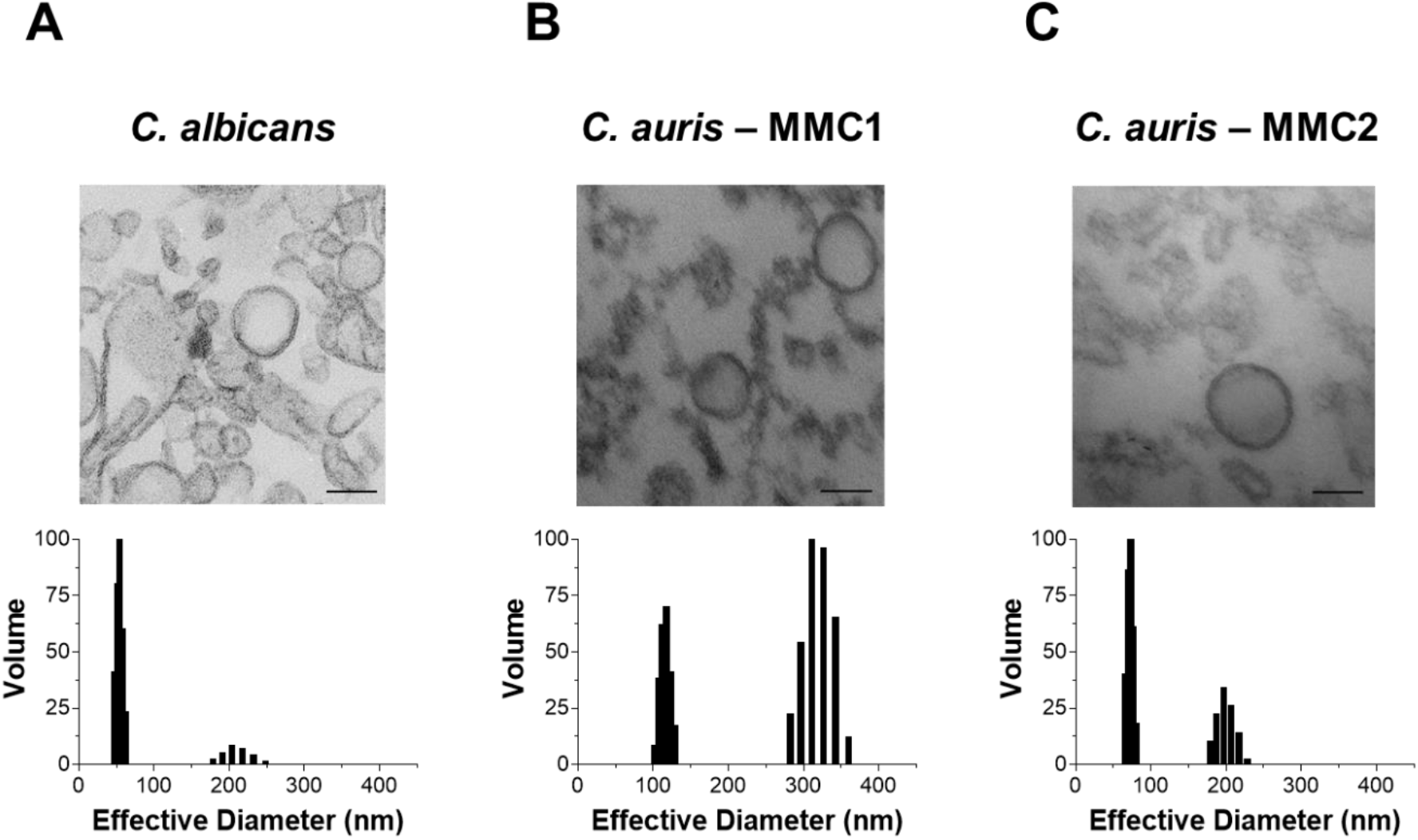
*C. auris* releases EVs. Transmission electron micrographs and dynamic light scattering measurements of EVs from *C. albicans* (A) and *C. auris* isolates MMC1 (B) and MMC2 (C). Two independent EVs isolations were analyzed by both methods with similar results. The figure shows representative results of each analysis. Scale bars = 100 nm

### Protein and ergosterol content of *C. auris* extracellular vesicles

The amount of protein (Fig. 2A) and ergosterol (Fig. 2B) were determined and normalized by the number of yeast cells in culture at the EV harvest time. Both *C. auris* strains secreted similar amounts of protein in EVs, but the amounts were 3-4 times lower compared to *C. albicans.* Likewise, the amounts of EVs ergosterol was 3-6 times lower in *C. auris* strains than in *C. albicans. C. auris* MMC2 strain EVs had the lowest amount of ergosterol, being 3 times lower compared to MMC1 (Figure 2B). As ergosterol is a ubiquitous molecule present in EVs membranes, we normalized the protein content by the amount of ergosterol in each strain as a way to measure possible differential protein loads among isolates. MMC2 had a higher protein/ergosterol ratio than either MMC1 or *C. albicans*.

**Figure 2.**
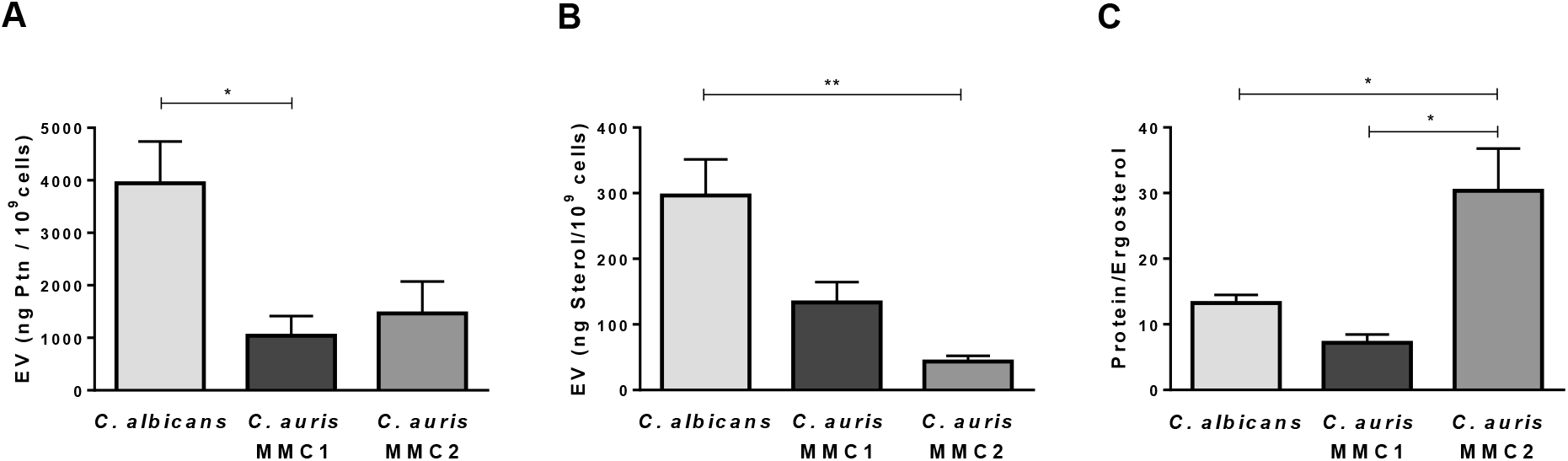
Protein and sterol content in *C. auris* and *C. albicans* extracellular vesicles (EVs). Protein (A) and sterol (B) concentrations were measured in EVs suspensions from *C. auris* and *C. albicans* and normalized by the number of cells present in the fungal cultures at the harvest time. (C) Protein to ergosterol concentration ratios. All graphs represent means and standard error of the mean, relative to 4 independent EVs isolations. *p ≤ 0.05; **p ≤ 0.01 by One-way ANOVA followed by Tukey’s multiple comparison test.

### RNA-seq analysis of *C. auris* extracellular vesicles

We performed next-generation sequencing from both MMC2 cells and vesicles to investigate their RNA profiles. This allowed not only to profile the EVs RNAs, but also to determine subpopulations of RNA that are enriched in the EVs compared to cells. The RNA-seq analysis led to the identification of 563 mRNAs and 104 ncRNAs in MMC2 EVs (Supplemental Table S1). From the transcripts identified in the EVs, the most enriched gene ontology terms were filamentous growth, transcription factor, and response to stress (Table 1).

**Table 1.** Gene ontology for RNAs enriched in EVs from MMC2

| Gene ontology Term - Extracellular Vesicles | Count | Fold Change | p-value |
| --- | --- | --- | --- |
| Biosynthetic process | 4 | 2.6 | 1.90E-02 |
| Cellular response to stress | 10 | 1.2 | 4.60E-02 |
| DNA replication, recombination, and repair | 4 | 1.5 | 4.90E-02 |
| Filamentous growth | 23 | 1.3 | 2.10E-02 |
| Growth of unicellular organism as a thread of attached cells | 3 | 2.9 | 2.70E-02 |
| Intracellular protein transport | 7 | 2 | 1.30E-02 |
| Transcription, DNA-templated | 16 | 1.2 | 3.20E-02 |
| Vesicle-mediated transport | 4 | 1.3 | 5.80E-02 |

| Gene Ontology Term - Cell | Count | Fold Change | P-Value |
| --- | --- | --- | --- |
| Carbon metabolism | 45 | 2.6 | 8.4E-10 |
| Translation | 35 | 3.0 | 5.0E-9 |
| Yeast-form cell wall | 21 | 5.1 | 1.7E-9 |
| Glycolysis / Gluconeogenesis | 24 | 3.2 | 5.4E-7 |
| Citrate cycle (TCA cycle) | 19 | 4.0 | 2.7E-7 |
| Tricarboxylic acid cycle | 13 | 5.2 | 2.7E-6 |
| ATP synthesis coupled proton transport | 6 | 4.4 | 9.1E-3 |
| Oxidative phosphorylation | 23 | 1.9 | 3.3E-3 |

When we compared the EVs and cellular mRNAs, the most abundant transcripts were different from each other (Figure 3A and 3B). The topmost abundant mRNAs in EVs code for uncharacterized proteins (QG37_06847) that present a domain of a peroxisomal membrane anchor protein (Pex14p). The second most abundant transcript is also an uncharacterized protein (QG37_03488) with an RNase III domain (Figure 3B). Indeed, most of the transcripts found in EVs are related to the metabolic process of RNA followed by regulation of biological process and organelle organization, although other functions are also present, as a response to both stress and drugs (Fig. 3B). Most of the ncRNAs were tRNA and tRNA-halves, followed by several small nucleolar RNAs and ribosomal RNAs (Fig. 3C). On the other hand, the most expressed mRNAs in cells were not the most represented in the EVs and they code for alcohol dehydrogenase 2, fructose-bisphosphate aldolase, and glyceraldehyde-3-phosphate dehydrogenase (Figure 3A). The most abundant pathways in cells were translation, cell wall, glycolysis/gluconeogenesis, and tricarboxylic acid cycle (Table 1). Overall, our results showed that *C. auris* EVs carry RNA, as demonstrated for other fungi ^45–47^. In addition, we showed that *C. auris* EVs have a different cargo compared to the whole cells, suggesting a selective process to load the transcripts to the EVs.

**Figure 3.**
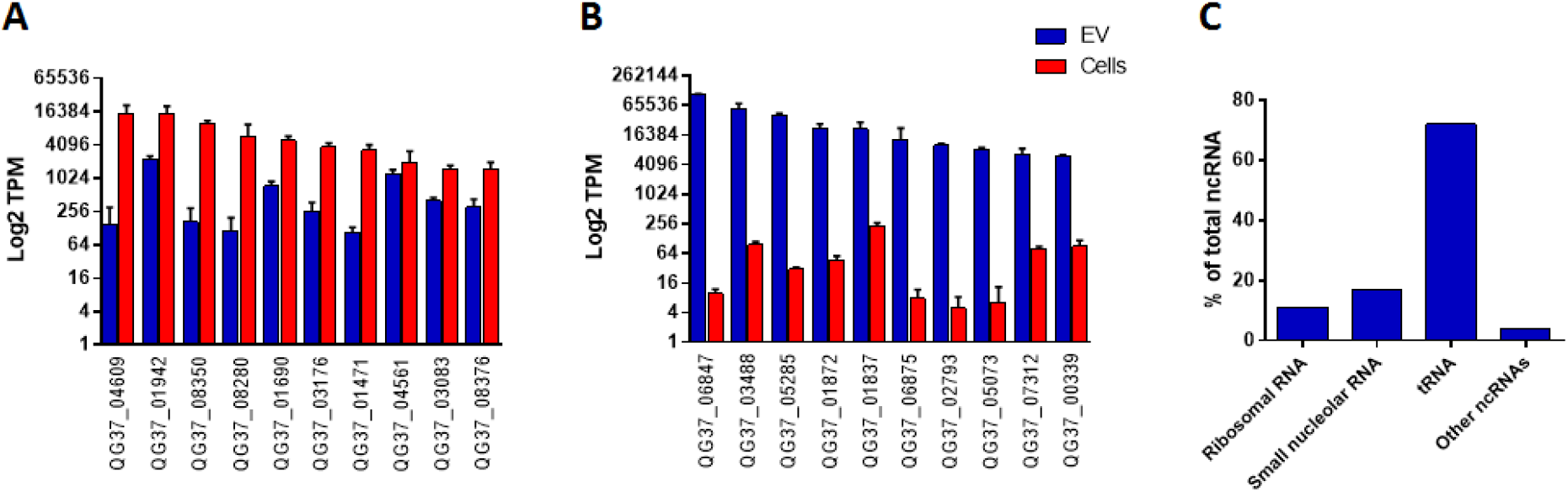
EVs from *C. auris* carry RNA. RNA was extracted from *C. auris* MMC2 yeast cells and EVs. Enrichment of RNAs was compared between yeast cells and EVs (6A and 6B). In C, the frequency for each class of non-coding RNA found in EVs from *C. auris* MMC2. EVs samples correspond to three independent EVs isolations.

### Proteomics analysis of extracellular vesicles

We performed EVs proteomic analysis and compared the protein profiles across both species, with a threshold of 40% of homology between their orthologues (Table S2). To make the comparison fair, we calculated the relative copy number of proteins per sample (% from total). We also compared the EVs data with the proteomics analysis of whole cells^39^, which were prepared and run in parallel. We observed striking differences between the whole cells and EVs for each of the 3 strains (Figure 4A). We performed hierarchical clustering to separate groups of proteins based on their abundance profile. Cluster 1, which contains proteins commonly enriched in EVs from *C. auris* and C. *albicans*, were enriched in proteins from starch and sucrose metabolism, protein processing in the endoplasmic reticulum, MAP kinases, and amino sugar and nucleotide sugar metabolism (Figure 4A). *C. albicans,* but not *C. auris* EVs, were enriched in abundant cellular proteins, such as ribosomal proteins and proteins from the central carbon and amino acid metabolisms (Cluster 2 in Figure 4A). EVs from both species were depleted of proteins from functions such as ribosomal biogenesis, proteasome, DNA replication, RNA degradation, and sterol biosynthesis (Clusters 3-5 in Figure 4A). Within the ten most abundant proteins in whole cells, only enolase 1 and pyruvate decarboxylase had high amounts in EVs (*C. albicans)* (Figure 4B). None of the top 10 most abundant whole-cell proteins were abundant in *C. auris* EVs (Figure 4B). On the other hand, the top 10 most abundant EVs proteins were present only in small amounts in whole cells (Figure 4B), suggesting a highly selective process to upload proteins into EVs.

**Figure 4.**
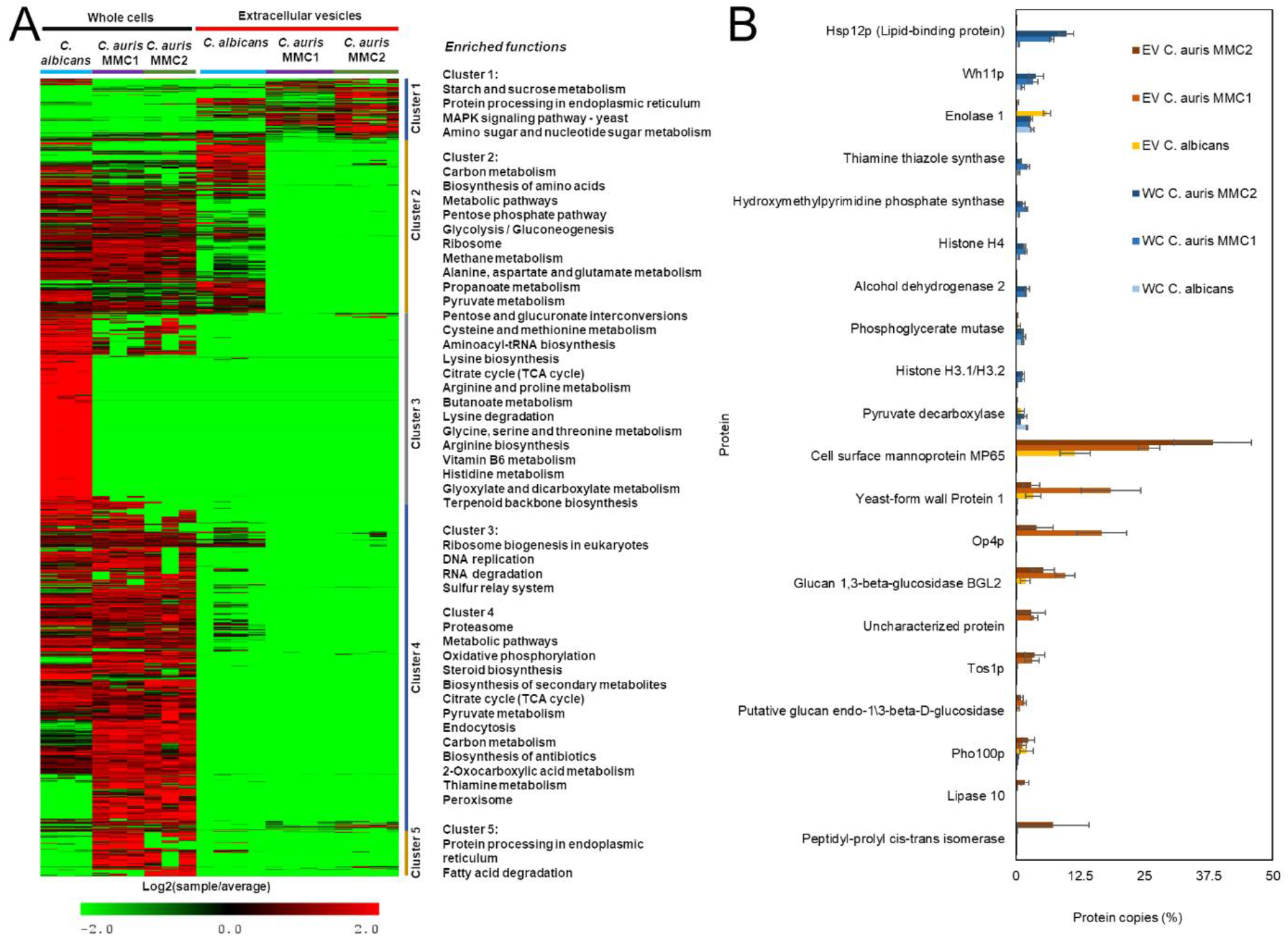
Comparative proteomic analysis of extracellular vesicles and whole cells from *C. albicans* and *C. auris*. Proteomic data of whole cells and extracellular vesicles were submitted to intensity-based absolute quantification (iBAQ) and converted to relative copy numbers before comparing across different samples. (A) Heatmaps and hierarchical clustering were performed with MeV and function enrichment of each cluster was performed with DAVID. (B) Profiles of the 10 most abundant whole-cell (WC) and extracellular vesicles (EVs) proteins. Cell samples correspond to three independent cultures and EVs samples correspond to 4 independent EVs isolations.

Among all the proteins detected in EVs, 393 were considered differentially abundant when comparing *C. auris* (both strains) with *C. albicans.* The number of differentially abundant proteins corresponded to 33% of the detected proteins, so the abundance of the remaining 66% was similar among the species. The heatmap in figure 5A shows all the major differentially abundant EVs proteins among the evaluated organisms. The heatmap was divided into three clusters based on differences of protein abundance between EVs from *C. auris* and *C. albicans.* Out of the 393 proteins on the heatmap, 42 proteins (~10%) were more abundant in *C. auris* than *C. albicans* (Cluster 2, Figure 5). This group of proteins was enriched in proteins from the starch and sucrose metabolism and protein processing in the endoplasmic reticulum (Figure 5B). As mentioned above, *C. albicans* EVs had higher amounts of metabolic proteins (Cluster 3, Figure 5A-B). *C. albicans* EVs had higher amounts of TCA cycle proteins (figure 5B), which is the opposite of what was found in whole cells ^39^. This result further supports the presence of a selective mechanism for sorting proteins into the EVs.

**Figure 5.**
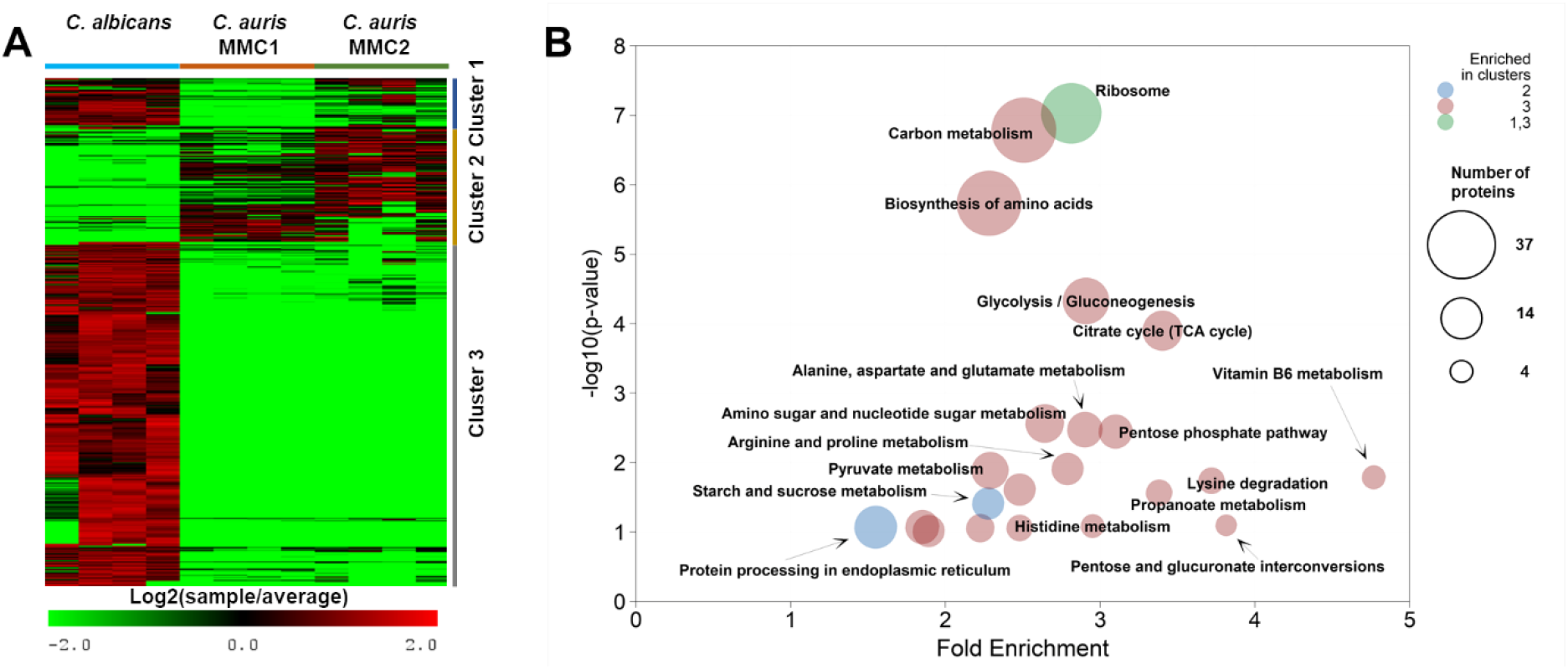
Proteomics analysis of EVs from *C. auris.* EVs from both *Candida* species were submitted to protein extraction and analysis. The heatmap (A) shows the abundance of proteins differentially abundant in EVs from both species of *Candida.* The heatmap was clustered using the hierarchical method. (B) Function-enrichment analysis of different clusters of proteins from panel A. The enrichment of pathways was done with DAVID. The graph represents the relationship between p-values and fold enrichment. The colors of the circles represent the different clusters they are enriched in, while the size, the number of proteins from each pathway. EVs samples correspond to 4 independent EVs isolations.

### Lipidomics analysis of extracellular vesicles

We performed a lipidomics analysis to compare the lipid profile of *C. auris* and *C. albicans* EVs (Figure 6A-B). All detected species of diacylglycerols (DG) and triacylglycerols (TG) were more abundant in EVs from *C. albicans,* whereas the majority of glycerophospholipids were enriched in the *C. auris* isolates, including phosphatidylcholine (PC), phosphatidylethanolamine (PE), phosphatidylglycerol (PG) and phosphatidylserine (PS) species (Figure 6A). Conversely, phosphatidic acid (PA) and phosphatidylinositol (PI) species were more abundant in *C. albicans* (Figure 6A). The pattern of sphingolipids was also distinct when the *Candida* EVs were compared. Two major species of conserved hexosylceramides (HexCer) were found in EVs from *C. albicans* and *C. auris* (Figure 6B), corresponding to the same distribution characterized in their respective yeast extracts recently reported by our group ^5^. Consistent with that, HexCer species bearing Cer(d18:1/24:0(2OH)) and Cer(d20:0/18:0) were more abundant in *C. albicans* EVs. Nonacylated sphingoid bases sphinganine Cer(d18:0/0:0) and sphingosine Cer(d18:1/0:0) were more abundant in *C. auris* MMC2 (Table S3). We also found unusual free ceramide species with acetate as the acyl group (Figure 6C), known as C2-ceramides, which were more abundant in *C. auris* MMC1. Remarkably, Cer(d18:1/2:0) comprises 7.9% of the mass spectrometry signal for all identified lipids in the positive ion mode analysis of *C. auris* MMC1 EVs, but only 0.6% and 0.2% of the *C. auris* MMC2 and *C. albicans* EVs, respectively (Figure 6C). We compared the relative intensities of Cer(d18:1/2:0) to the whole cell data from our recent publication^5^. We found an enrichment of 25, 67, and 109 folds of this lipid species in EVs compared to the whole cells in *C. auris* MMC1, *C. auris* MMC2 and *C. albicans*, respectively.

**Figure 6.**
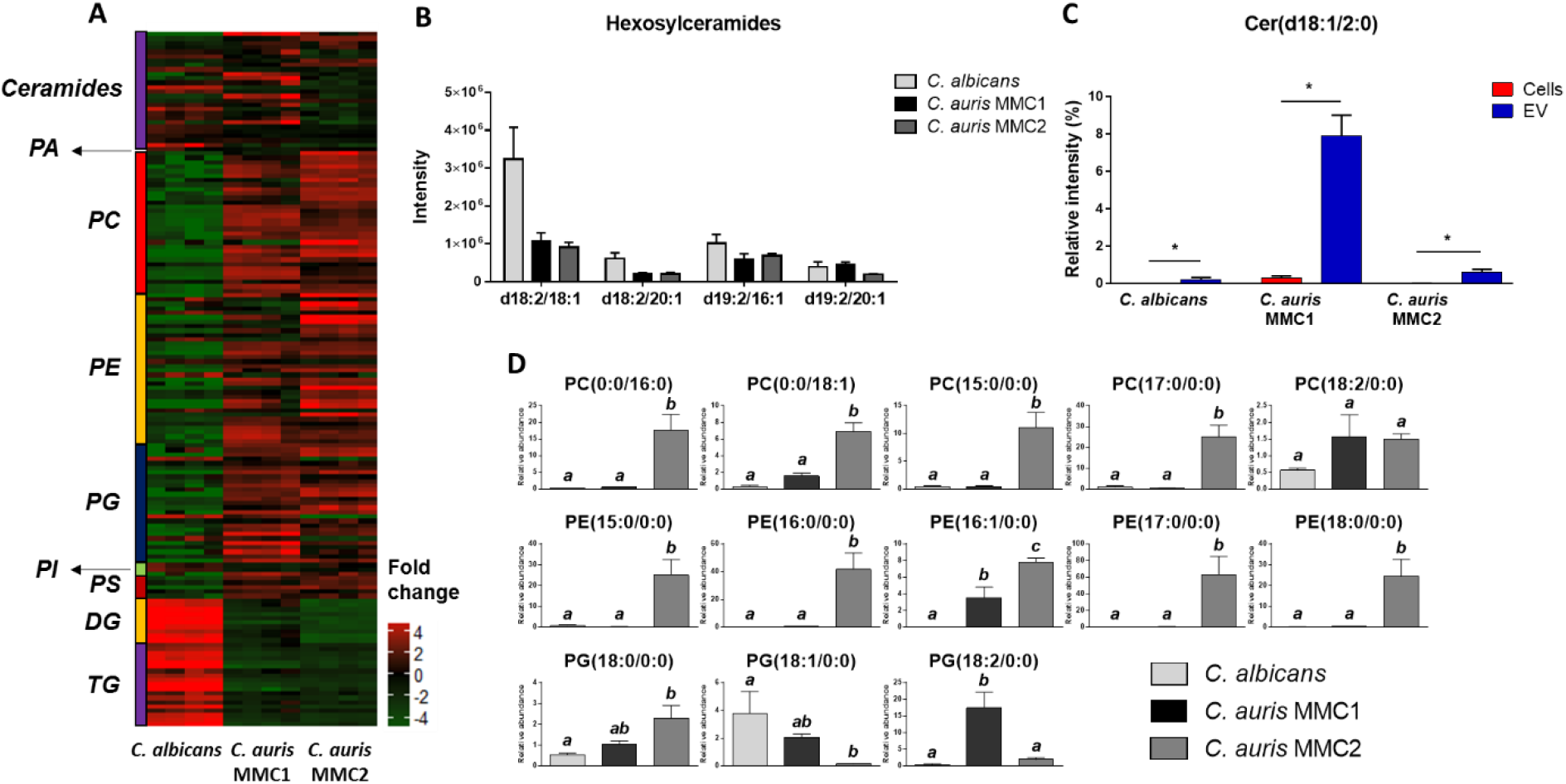
Lipid profile of *C. auris* and *C. albicans* EVs. Vesicles from both *Candida* species were submitted to lipid extraction and analysis by LC-MS/MS. (A) Heatmap of the relative abundances of EVs lipids from both *Candida* species. (B) Relative intensity of hexosylceramides. (C) Relative intensity of Cer(d18:1/2:0) was compared between EVs and yeast cells among all Candida species. (D) Relative abundance of lysophospholipids In *C. auris* and *C. albicans* EVs. Data were analyzed by one-way ANOVA followed by Tukey’s multiple comparison test. Different letters among bars represent *p* < 0.05. EVs samples correspond to 4 independent EVs isolations. Abbreviations: Cer, ceramide; DG, diacylglycerol; PA, phosphatidic acid; PC, phosphatidylcholine; PE, phosphatidylethanolamine; PG, phosphatidylglycerol; PI, phosphatidylinositol; PS, phosphatidylserine; TG, triacylglycerol.

Recently, we reported that *C. auris* had higher expression of a variety of phospholipases compared to *C. albicans* ^5^, therefore, we took a closer look at their products, lysophospholipids^48^. *C. auris* MMC2 EVs had a consistent higher abundance of lysophosphatidylcholine (LPC) and lysophosphatidylethanolamine (LPE) species compared to *C. auris* MMC1 and *C. albicans* (Figure 6D). Lysophosphatidylglycerol (LPG) species, being PG(18:0/0:0), PG(18:1/0:0), and PG(18:2/0:0) more abundant in *C. auris* MMC2, *C. albicans* and *C. auris* MMC1, respectively (Figure 6D).

Fatty acids (FA) chains were detected in all organisms and ranged in size from 14 to 24 carbons, and arachidonic acid was detected, esterified to phosphatidylethanolamine (PE(18:2/20:4)), consistently in both *C. auris* isolates. Arachidonic acid was not abundant in the evaluated strain of *C. albicans* in the tested conditions (Table S3).

### Effects of extracellular vesicles on the yeast adhesion to epithelial cells

Adhesion to epithelial surfaces is an important feature displayed by pathogenic species of *Candida* as an early stage of colonization of host tissues ^49–52^. We evaluated whether *C. albicans* or *C. auris* EVs had an impact on the adhesion of *C. auris* or *C. albicans* to HeLa epithelial cells monolayers. Pre-incubation of HeLa cells with *C. auris* MMC1 EVs increased the adhesion of this yeast. The same was not seen for MMC2 or for *C. albicans,* as the pre-treatment with their EVs did not affect the adhesion of yeast to HeLa cells (Figure 7). This result shows how EVs from different strains of the same species can induce distinct host cells phenotypes. The graph shows average and standard errors relative to 2 independent experiments. \**p* < 0.05 by one-way ANOVA followed by Dunnet test.

**Figure 7.**
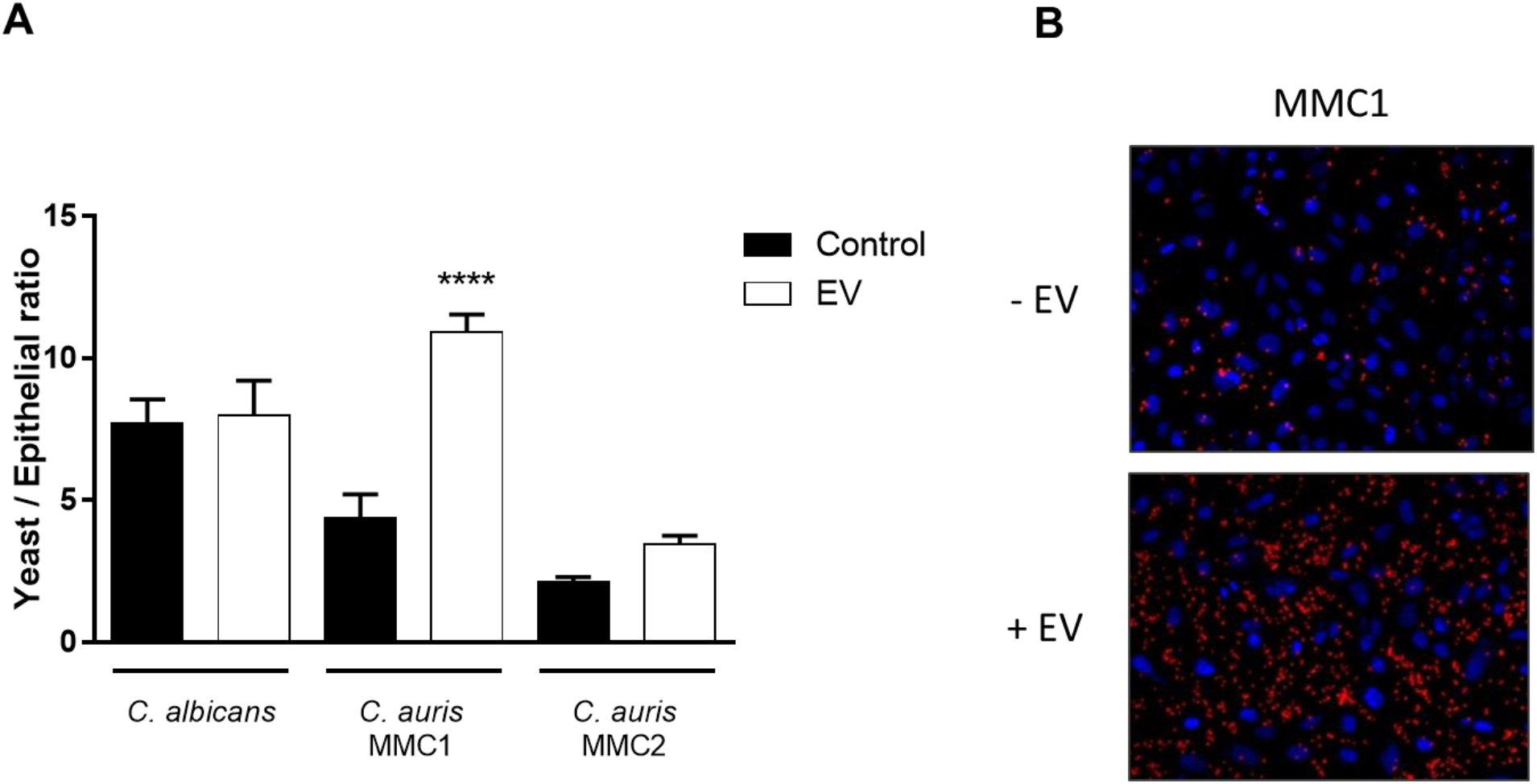
The effect of extracellular vesicle (EVs) pretreatment on the adhesion of yeast cells to epithelial monolayers. HeLa cells were pre-treated, or not, with EVs (10 μg/mL of protein) for 1 h before challenging with the respective yeast cells. After incubating for 1 h, monolayers were washed, and slides were analyzed under a fluorescence microscope (see Methods for details). (A) Quantification of adhering yeast cells. (B) Fluorescence image of *C. auris* MMC1 adhesion to HeLa cells. Nuclei were stained with DAPI (blue), whereas the yeasts were stained with NHS-Rhodamine (red).

### Effect of extracellular vesicles on phagocytosis and killing by macrophages

EVs from certain fungi can affect the way yeast cells are internalized and killed by macrophages ^8,13–16,53^. We tested whether EVs from *C. auris* or *C. albicans* would be able to modulate the uptake or clearance of yeast cells by macrophages. The incubation with EVs from either *C. albicans* or *C. auris* had no significant effect on the phagocytosis of yeast cells by macrophages (Figure 8A). However, whereas EVs pre-incubation enhanced macrophages’ ability to kill *C. albicans*, EVs from *C. auris* MMC2 but not MMC1, enhanced yeast cell proliferation within the macrophages (Figure 8B). This points to differing roles for EVs among species regarding some of the effector functions of macrophages.

**Figure 8.**
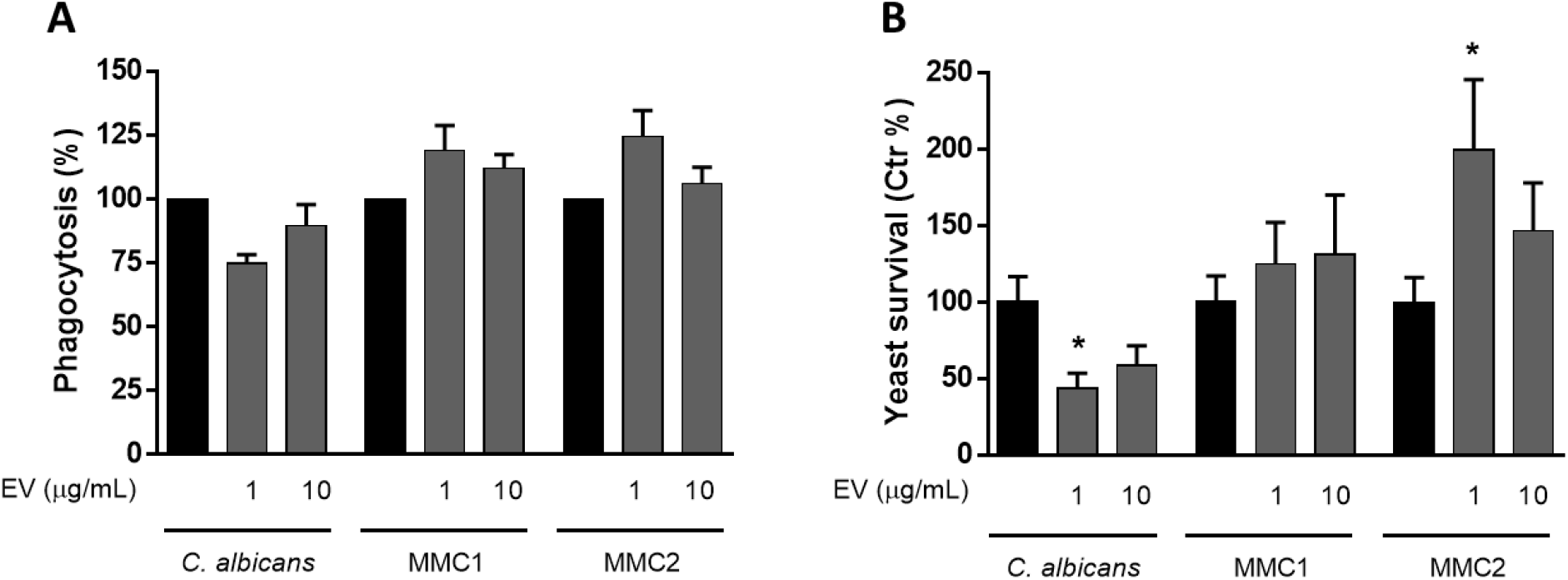
Phagocytosis and killing by macrophages. (A) RAW 264.7 macrophages were incubated with EVs for 1 h and then challenged with yeast cells in the ratio of 1:2 (macrophage:yeast) for 1 h. After this period, extracellular yeast cells were washed off and macrophages were lysed and plated onto Sabouraud for CFU counting. (B) Bone marrow-derived macrophages were incubated with EVs for 4 h and then challenged with yeast in the ratio of 10:1 (macrophages:yeast) for 24 h. The macrophages were then lysed and plated onto Sabouraud for CFU counting. Graphs show averages and standard error on the mean for 4 independent experiments. *\*p* < 0.05 by paired T-test.

### Activation of dendritic cells by extracellular vesicles

We investigated the ability of *C. auris* to regulate dendritic cells by measuring 3 important signals for antigen presentation and activation of T cells, MHC-II, co-stimulatory molecules (CD80 and CD86), and cytokines ^54^. BMDC were incubated for 24 h with *C. auris* or *C. albicans* EVs, and MHC-II and co-stimulatory molecules were measured by FACS, whereas cytokines were assayed by ELISA. We observed an increase of surface markers associated with BMDC activation, which was similar to the one induced by LPS. Despite that, MMC2 EVs induced a lower response, and all tested EVs concentrations from *C. auris* and *C. albicans* were able to increase, in a dose-dependent manner, the expression of MHCII, CD80, and CD86 on BMDC (Figure 9A-C). BMDC treated with EVs from both *Candida* species did not produce IL-10, IL-12 or TNF-α. However, BMDC stimulated with EVs from *C. auris* produced IL-6 similarly to *C. albicans* (Figure 9D). In addition, inhibition of the basal production of TGF-β by BMDC was detected after the incubation with EVs from *C. auris* MMC1 and *C. albicans* (Figure 9E).

**Figure 9.**
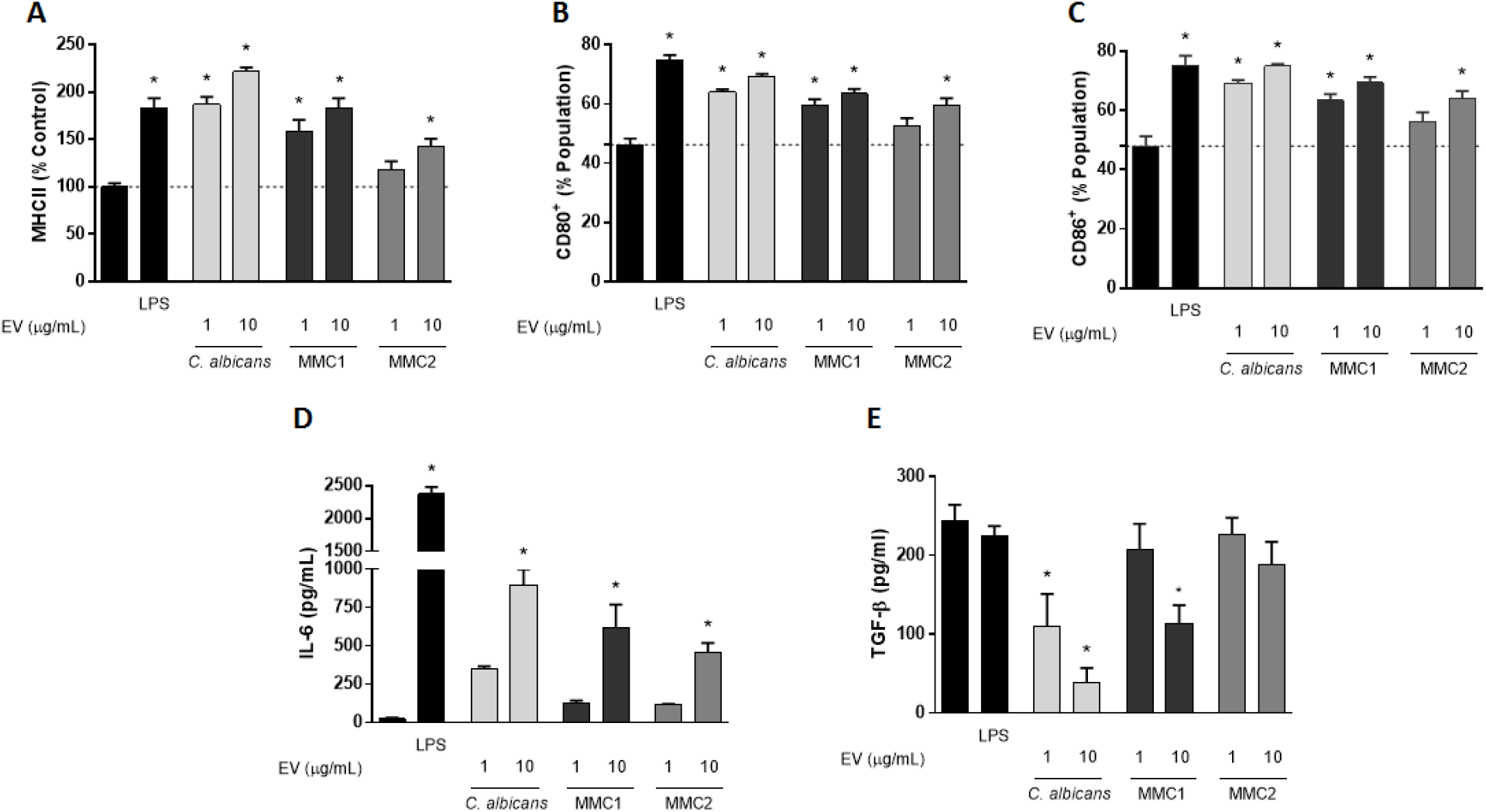
Activation of BMDC by *C. auris* extracellular vesicles (EVs). BMDC were incubated, or not with EVs from *C. albicans* and *C. auris* for 24 h and then analyzed by flow cytometry (A-C) for the expression of MHC-II (A), CD80 (B) and CD86, or ELISA (7D and 7E), for the production of IL-6 (D) and TGF-β (E). Graphs show mean and standard error for 3 independent experiments. * = p < 0.05 by ANOVA followed by Tukey.

## Discussion

The literature reports two distinct populations of EVs from other species of *Candida,* a smaller one ranging usually from 50-70 nm, and a larger group between 100 and 800 nm 9,25,^44^. We found that *C. auris* releases EVs encompassed by lipid bilayers with size and shape consistent with those from other species. TEM showed that, as opposed to some species such as *Histoplasma capsulatum* ^25^ and *Cryptococcus neoformans* ^24^, *C. auris* EVs lack electron-dense areas associated with pigmentation. The total content of protein and ergosterol in EVs suspensions was higher in *C. albicans* than in either *C. auris* strain when normalized by the number of producing cells. However, *C. auris* contained more protein and ergosterol in EVs suspensions than *C. albicans* (data not shown), probably due to the higher cell count in *C. auris* cultures. The ratio between proteins and sterol excludes the number of cells from the analysis and *C. auris* MMC2 had a ratio higher than *C. albicans* or *C. auris* MMC1.

EVs RNAs have been characterized in *C. neoformans, S. cerevisiae, P. brasiliensis, H. capsulatum* and *C. albicans* ^45,46^. We found that the most abundant transcripts in *C. auris* EVs are different from the whole cells, suggesting a selective packing for RNAs during EVs biogenesis. The most abundant transcripts of EVs from *C. auris* MMC2 were associated with ribonuclease, GTPase and ubiquitin activities or from uncharacterized genes. EVs mRNAs of *C. albicans* and *C. auris* share common biological processes, such as cellular response to stress and filamentous growth, indicating a conserved sorting mechanism ^45^. In other eukaryotes, EVs mRNAs can be translated into the recipient cell ^55^, although this should be experimentally addressed for fungal EVs. The most abundant ncRNAs in EVs from *C. auris* MMC2 were tRNAs and their fragments, similar to previously described for *C. albicans* (~60%). These fragments of tRNA have been described in EVs of diverse organisms, from unicellular parasites to human cells ^56–58^. In T lymphocytes-derived EVs, the most abundant class of RNA characterized is tRNA fragments, comprising 45% of all RNA identified in the EVs compared to the cell content, and these fragments act by repressing immune activation in T cells ^59^.

The EVs proteomic profile was strikingly different from the cells they are derived from. Whereas *C. auris* EVs were enriched in proteins from the starch and sucrose metabolism and protein processing in the endoplasmic reticulum, *C. albicans* EVs had higher amounts of proteins from the central carbon metabolism, ribosomes, and amino acid metabolism. We have previously shown that TCA cycle proteins were more abundant in *C. auris* yeast cells than *C. albicans* ^39^, but we are now showing an opposite phenotype in EVs, suggesting a selective sorting that could help control the intracellular levels of specific metabolic enzymes. These differences suggest that the EVs from *C. albicans* and *C. auris* are involved with distinct metabolic adaptations. In terms of lipid composition, the relative abundance of lipids involved with energy storage, as triacyl- and diacylglycerols (TG and DG), is remarkably higher in EVs from *C. albicans,* when compared to the *C. auris* ones, reflecting the pattern found in their originating yeast cells ^5^. The relative abundance of structural glycerophospholipids is consistently higher in EVs from both isolates of *C. auris,* also reflecting the lipid profile of their generating yeast cells ^5^. Although for some cases the lipid profile from *C. auris* EVs resembled the yeast cell one, some lipids from the yeast cells were not present in EVs, such as cardiolipins, which are mitochondrial markers. The amount of HexCer correlated with the distribution in their respective cells, as previously showed by our group ^5^. Considered initially as membrane structural components, HexCer were described as virulence regulators in *C. albicans* and *C. neoformans* ^60,61^. Their role in EVs could be linked to membrane and lipid raft stability ^62^. However, recently, Xisto and colleagues demonstrated that purified HexCer produced by the opportunistic fungus *Lomentospora prolificans* induced an oxidative burst by and increased the antifungal activity of macrophages ^63^.

To our knowledge, our findings report the first time that a C2-ceramide derivative has been found in fungal EVs. In mammalian models, C2-ceramide has biological properties such as antitumoral activity inducing apoptosis and arresting cell cycle^64,65^. The relative abundance of lysophospholipids was considerably higher in EVs from *C. auris* than *C. albicans,* particularly in EVs from MMC2. This data suggests an intense activity of lipid catabolic enzymes in *C. auris,* such as phospholipases. Some lysophospholipids are biologically active on leukocytes, for instance, LPC released by apoptotic neutrophils recruit monocytes from the bloodstream to promote clearance of apoptotic bodies from tissues ^66^. Immunomodulatory properties of LPC were demonstrated for other infection models and LPC could act as a virulence factor in *C. auris* infections ^67,68^.

To examine the potential biological effects of EVs upon host cells and to compare biological activities between species, we used amounts of EVs based on protein concentration in the same range as used in other reports for host-pathogen studies ^8,9,16,17^. Within this concentration range, EVs from other pathogenic fungi, were proven to be biologically active in distinct models. Adhesion of yeast cells to epithelial surfaces is an important mechanism of disease as an initial step for further tissue damage and colonization of distinct sites in the host, including the bloodstream ^52^. *C. albicans* can interact with surface adhesion molecules on epithelial cells ^50,51^, so we investigated whether EVs from both *Candida* species were able to modulate the adhesion of yeast to epithelial monolayers *in vitro.* Although the tested strains were able to adhere to the epithelial monolayer, a significant increase in adhesion was observed when EVs from *C. auris* MMC1 were added to the monolayer. Notably, *C. neoformans* EVs fuse with brain microvascular endothelial cells, changing their permeability ^22^ Since in our studies the epithelial cells were incubated with EVs prior to the challenge with yeast cells, it is possible that fusion with the epithelial cells could modify their permeability and/or modulate the exposure of adhesion molecules, although further experimentation is needed to address this hypothesis. Molecules involved with the adhesion of *C. albicans* to epithelial cells have been described, such as *C. albicans* ALs3p and Eap1p, and are potential players in the increase of adhesion induced by EVs ^69^.

Fungal EVs can induce the activation of phagocytes, increasing phagocytosis, cytokine production, and antigen presentation ^8–10,17^. EVs isolated from both *C. auris* strains did not modulate the uptake of yeast cells by macrophage cell lines as RAW, but EVs from *C. auris* MMC2 inhibited the killing of yeast cells by BMDM. EVs from *C. albicans* increased the killing of yeast cells by BMDM. EVs from other pathogenic fungi can modulate phagocytosis and/or killing by macrophages, but our data shows that EVs from only one of the *C. auris* isolates (MMC2) could inhibit the killing of the pathogen by macrophages. This data suggests that EVs from the same species could promote distinct changes in host cells. EVs from *C. albicans* followed the pattern played by most fungal EVs, as they induced killing ^8,10,14–16,53^. The incubation with EVs stimulated BMDC to express important signals responsible for CD4+ T cells activation such as MHCII, CD80 and CD86. The secretion of TNF-α, IL-10 and IL-12p70 by BMDC was not detected at biologically relevant levels. However, EVs from both *C. albicans* and *C. auris* induced the release of IL-6 by BMDC, while decreasing the basal production of TGF-β. This suggests that EVs from *C. albicans* and *C. auris* MMC1 induce an inflammatory response in BMDC. Apart from previously reported ^9^, TNF-α, IL-10 and IL-12p70 were not produced by BMDC stimulated with EVs. Different strain of *C. albicans* were used these studies, reinforcing the possibility that the biological activity of EVs could be strain-specific.

In summary, our results show that the emerging pathogen *C. auris* produces EVs that are similar in size to other pathogenic fungi, but the content of these EVs distinctly differs from what is known for *C. albicans* and these differences could explain the phenotypic changes induced by these EVs in the cells from the host. In this regard, we note that *C. auris* is a new fungal pathogen that has been proposed to have emerged from the environment as a result of global warming ^70^. In contrast, *C. albicans* has an ancient association with human hosts. Thus, the similarities in structure and content between *C. auris* and *C. albicans* EVs probably reflect constraints common to fungal cells and their physiology, while the differences reflect species-specific variables and perhaps the result of differences in the time of adaptation to human hosts.

## Acknowledgements

The Johns Hopkins University School of Medicine Microscope Facility. D.Z.M., J.D.N. and E.S.N. were supported by National Institute of Health-National Institute of Allergy and Infectious Diseases R21 AI124797. L.N. was supported by grants from the Brazilian agency Conselho Nacional de Desenvolvimento Científico e Tecnológico (CNPq, grants 311179/2017-7 and 408711/2017-7) and FAPERJ (E-26/202.809/2018). M.L.R. was supported by grants from the Brazilian Ministry of Health (grant number 440015/2018-9), Conselho Nacional de Desenvolvimento Científico e Tecnológico (CNPq, grants 405520/2018-2, and 301304/2017-3) and Fiocruz (grants PROEP-ICC 442186/2019-3, VPPCB-007-FIO-18 and VPPIS-001-FIO18). M.L.R. also acknowledges support from the Instituto Nacional de Ciência e Tecnologia de Inovação em Doenças de Populações Negligenciadas (INCT-IDPN). A.C. was supported in part by NIH grants AI052733, AI15207 and HL059842. Parts of this work were performed in the Environmental Molecular Science Laboratory, a U.S. Department of Energy (DOE) national scientific user facility at PNNL in Richland, WA.

## Notes

### Competing Interest Statement

The authors have declared no competing interest.

